# Aggressive interactions between smooth-coated otters and water monitor lizards in Singapore

**DOI:** 10.1101/2022.07.01.498521

**Authors:** Haaken Zhong Bungum, Philip Johns

## Abstract

Smooth-coated otters (*Lutrogale perspicillata*) and Malayan water monitor lizards (*Varanus salvator*) occupy similar habitats and and interact regularly in Singapore’s waterways. These interactions have a range of potential outcomes and are sometimes lethal. Few formal behavioral studies exist for either species. We analyzed interactions between otters and monitor lizards by gleaning data from publicly available videos from citizen scientists to examine what factors influence aggressive and defensive behaviors, and what influences vigilance in otters. Behavioral sequence analysis revealed no obvious monitor lizard behavior that predicted otter aggression towards monitors. We found that the presence and number of otter pups is positively associated with otter aggression. Otters also tended to be more vigilant in groups with more pups, and more vigilant on land than water. Monitor lizards displayed aggressive and defensive behaviors more frequently than did otters, regardless of whether the otters were aggressive towards lizards. These observations suggests that otters vary their aggression and vigilance levels depending on the context of each interaction.

## Introduction

Smooth-coated otters are medium-sized aquatic carnivores (about 10 kg) that live throughout South and Southeast Asia (see Hwang & Larivière 2005 for review). They live in both freshwater and brackish environments where they feed on fish and other aquatic organisms. Smooth-coated otters are cooperative breeders that live in family groups (“romps”) of up to 14 or more individuals (Hwang & Larivière 2005, Khoo & Sivasothi 2018a, 2018b, personal observation). They function as the apex predator in many ecosystems and can be a sign of healthy waterways (Theng et al. 2016). In Singapore, otters had disappeared by the 1970s, coinciding with increased urban development; partly due to an active campaign to clean Singapore’s waterways, smooth-coated otters have returned (Khoo & Sivasothi 2018b), and sightings of smooth-coated otters surged around 2014, in part because otter watchers in Singapore could easily share their sightings over social media, along with the rapid increase in the otter population (Theng & Sivasothi 2016). Singapore’s otters have been extremely successful in their recolonization of local waterways (Khoo & Lee 2020), and there are now at least 11 family groups across Singapore (Khoo & Sivasothi 2018b) as well as lone otters not attached to any romp.

Otters encounter other species, sometimes with negative consequences. Feral dogs can be aggressive to otters and vice versa (Clements 2019), and social media includes reports of otters harassing estuarine crocodiles (*Crocodylus porosus*; Toh 2018), a behavior similar to that of giant otters (*Pteronua brasiliensis*), which sometime attack caimans (Ribas et al. 2012). Smooth-coated otters also frequently encounter Malayan water monitor lizards (*Varanus salvator*). These large lizards (up to 20 kg) inhabit a range of tropical environments from the Molucca Islands to Sri Lanka (Twining & Koch 2018). Water monitors are opportunistic, and like smooth-coated otters they prey on fish and other animals, but unlike otters also frequently scavenge (Twining & Koch 2018). Monitors persisted in Singapore, even as otters were driven out by a deterioration of suitable habitat. Today, they are common in Singapore’s waterways, whether concrete canals or natural riverbanks.

Interactions between otters and monitors can take several forms. Water monitors can be commensalists that scavenge fish remains otters leave (personal observations), kleptoparasites that steal fish from otters (e.g., Tan 2019), competitors for food resources, or predators that attack young otters (e.g., Lee 2019; personal observations). Sometimes otters and monitors interact aggressively (Figure 1), which can lead to otters attacking monitors and to the injury or even death for the monitor lizard (e.g., Mitchell 2021). Otter-monitor conflict has been noted before (Goldthorpe et al. 2010), albeit from a single observation in Peninsular Malaysia.

**Figure 1.**
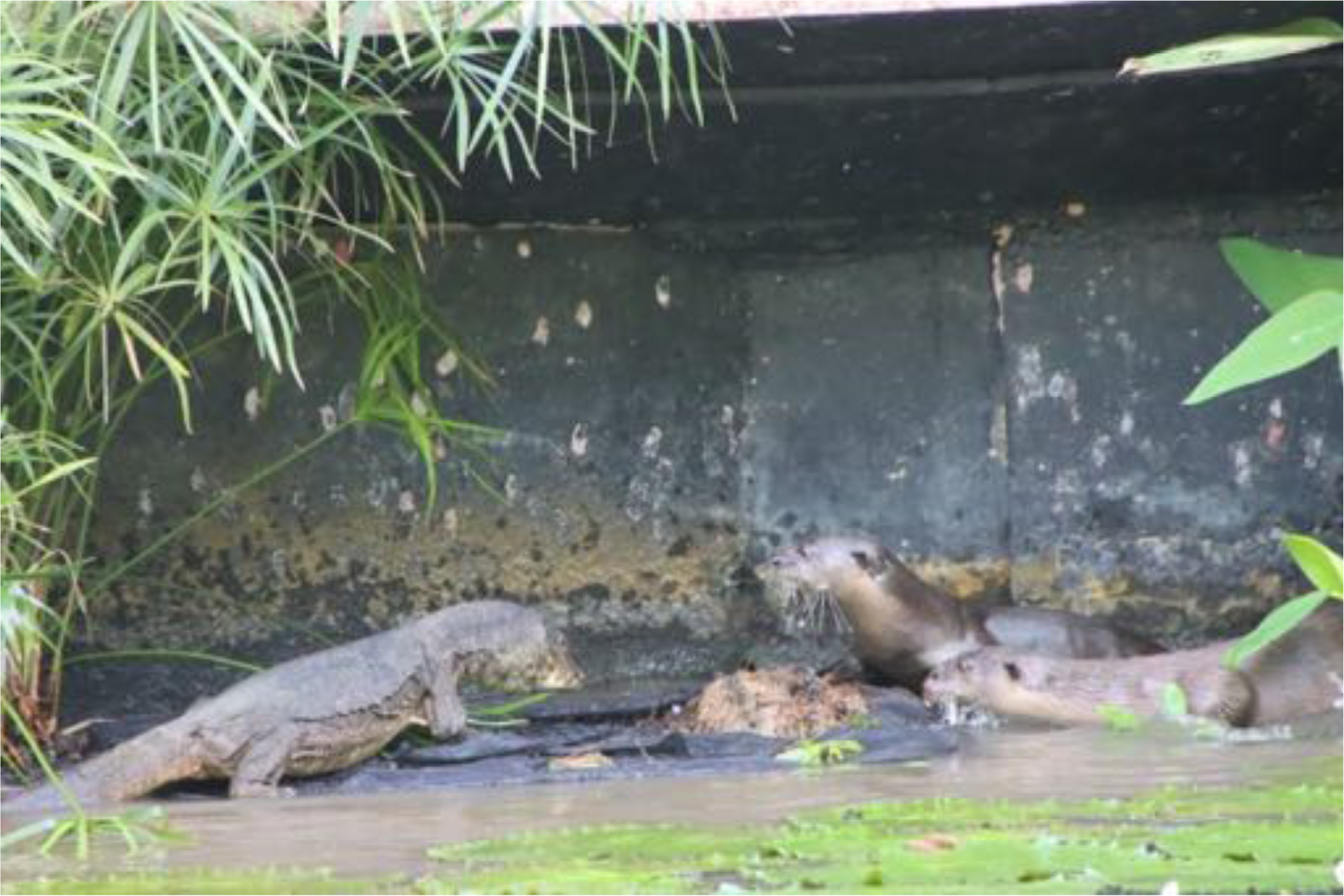
Two smooth-coated otters approach a monitor lizard in Singapore. Notice that the monitor lizard’s neck frill is extended in an aggressive manner. The monitor whipped its tail at the otters shortly after this photo was taken. Photo courtesy of Alicia Ellen Brierley.

Few formal studies have been conducted on either smooth-coated otter or monitor lizard behaviors. The demographics of smooth coated otters in Singapore and elsewhere are well-studied (e.g., Theng & Sivasothi 2016; Theng et al. 2016; Khoo & Sivasothi 2018a, 2018b), but there is scant literature on their behavioral traits, especially those that may be unique to urban otters. Urban environments change behaviors in other animals (e.g., Slabbekoorn & den Boer-Visser 2006; Breck et al. 2019), and urban otters also adjust to their environments. They construct unique holts in manmade structures (Khoo & Sivasothi 2018b) and use manmade features like ladders in their movements (Lay 2021). The robust population of otters in Singapore is notable because in other areas, smooth-coated otters are associated with natural landscapes (Kamjing et al. 2017). Likewise, outside of a few ecological and descriptive studies (e.g., Twining & Koch 2018; Uyeda 2009), very little is known about water monitor behaviors, especially in urban environments.

These sometimes-violent encounters between otters and monitor lizards raise questions of their causes. In other species, otter groups can be formidable. In one study, giant South American otters (*Pteronura brasiliensis*) mobbed jaguars until the jaguars left the area, but a lone otter did not engage the larger cat by itself (Leuchtenberger et al. 2016). Smooth-coated otters live in family groups (“romps”) like those of giant otters, and group dynamics may play a role in their interactions with water monitors. Otter pups are particularly vulnerable; the presence and number of pups may influence otters’ responses. Likewise, whether monitor lizards’ are aggressive or defensive may depend on both otter behaviors and the otter group composition. Otters are vigilant to potential threats, adopting a characteristic upright posture and other behaviors in and out of water, but the relationship between group size, the number of pups present, and how vigilant otters are to monitors or other threats, has not been addressed. Here we examine encounters between otters and monitor lizards to assess factors that lead to aggressive interactions, and specifically to otters attacking monitor lizards.

Collecting behavioral data can be extraordinarily time consuming, especially data related to uncommon events like otter-monitor interactions. However, the ease with which consumer-grade technology like smart phones and relatively inexpensive cameras can record photos and videos has led to an explosion of raw data on social media and other corners of the internet. Crowd-sourced data has been used for citizen science studies related to occupancy, ecology, and conservation for some time (see Cooper 2016 for review). Citizen science can also be effective in animal behavior studies, and some studies glean data from public sources (e.g., Boydston et al. 2018; Loong et al. 2021; Bungum et al. 2022). Gleaning data from public repositories, such as social media or YouTube, can be particularly useful when the behavior or species is rare or difficult to find (Nelson & Fijn 2013). Here we glean data from publicly available sources of video to address otter-monitor interactions. Because social media posts amount to *ad lib* sampling (Altmann 1974), there is a potential for bias. We try to avoid that by narrowing the scope of our study to otter-monitor interactions.

## Materials and Methods

Almost all data in this study came from online sources due in part to the restrictions imposed by COVID-19 pandemic. Selected videos were obtained from YouTube (www.YouTube.com) using search terms such as “Singapore otter monitor” and “Singapore otter lizard” (Appendix 1). One YouTube channel had a large collection of videos, “RandomSG” (Wong 2019; www.youtube.com/channel/UCLz7pIXxzaFzz_MN02kNzsQ), and the videos posted on this channel were not edited. Otters in Singapore’s waterways are generally active early and late in the day (roughly 06:30 to 10:00 and 16:00 to 19:30), and there was a nearly complete record of twice-daily videos spanning 2019-2021, ceasing only during the mandated lockdowns due to COVID-19. The regular recordings were free of obvious bias and allowed us to track pup age over time. We identified 160 videos with potential interactions between otters and monitors; due to quality, duplications, and scoring protocols, we scored 63 for the study, all taken between October 2018 and July 2019, of the Pandan otter romp, which resided near the Ulu Pandan River in southwestern Singapore. (In Singapore, people name otter romps after the first place they were observed; in 2019, after the events in this study, the Pandan romp was forced out of the river by the Jurong Lake Gardens romp.) The Pandan romp consisted of 9 adults and up to 9 pups at the time. Videos provided were taken using a handheld, fixed-lens DSLR camera, generally positioned at the bank opposite the otters’ activities, 10-30 m away.

These 63 videos were chosen from a time of 70 days before the first emergence of the pups in May 2019 until 40 days post-emergence. Pups remain in the holt for roughly six weeks after birth (Khoo and Sivasothi 2018a). At this age, pups are not yet fully weaned, and so are still heavily reliant on romp members’ help to survive (Hwang & Larivière 2005). This chronologically complete set of videos allowed us to compare behavior of adults before pups were born, after pups were born but before they emerged from their holt, and after the pups emerged and as they aged.

First we examined the association between otter and monitor aggression in all 63 videos without distinguishing between potential aggression based on a minimum distance criteria. If monitor lizards and otters came within one otter body length of each other (about 1m), that encounter had the potential for direct aggressive. Videos with the potential for aggressive encounters were then divided into “bouts”, where each bout lasted up to one minute before and after: 1) an aggressive behavior between otters and monitors; 2) the first instance of otters and monitors coming within one otter body length of each other; or 3) until the subjects left the frame and scoring ceased within the one-minute window. Videos without the potential for aggression could not be separated into bouts because otters and monitors were never close enough to physically interact, although monitors were free to display a defensive posture. (It is difficult to disentangle monitor aggression and defense, and here we group monitor aggressive and defensive behaviors.) We treated bouts within videos as independent events. The bout ended if the lizards and otters turned away and disengaged from each other; if they performed a “benign” rather than aggressive or defensive behavior – typically something like “sit” or “groom” – or if a focal otter interacted with another otter. By these criteria the average bout length was 53.24 + 35.78s (median 50.49s). Otters are extremely active and often engage in several behaviors per minute. In the one incident in which there were back-to-back bouts between the same monitor and otter, the behavior that ended one bout began the next. By these criteria we observed 234 bouts within the 63 videos. Each bout included 1 to 18 otters and f 1 to 3 lizards. All 234 bouts reached a conclusion before the videos cut off, and there were no recorded serious injuries in any of the scored observations. There were also no edits or other post-production cuts within videos, i.e., the videos accurately reflected interactions. (Many otter watchers in Singapore edit their videos before posting them.)

We analyzed behavioral sequences in the 234 bouts to describe typical behavioral exchanges between otters and monitor lizards, and to examine possible suites of behaviors that might spur attacks. In our behavioral sequence analysis, we included behaviors from both otters and monitors to address cause and effect between species. We used four pilot videos, excluded from the final analysis, to generate an ethogram of otter and monitor behaviors (Table 1; Figure 2). We scored the 63 videos using the Behavioral Observation Research Interactive Software (BORIS; (Friard & Gamba 2016; http://www.boris.unito.it), using an all-occurrences approach (Altmann’s 1974). Smooth-coated otters usually lack individual markings, which precluded our tracking individuals from one video to the next. Within each video, we assigned each otter and each monitor lizard an identifier (“subject”) based on the animal’s initial position at the start of the interaction. If either the otter or lizard exited the frame, we ceased all scoring, i.e., both otter and monitor needed to be in view for any scoring to occur. If both subjects returned to the frame, they retained their subject designation only if we could be certain they were the same individuals. If there were any doubt or lack of clarity, subjects were reassigned and we analyzed subsequent behaviors as a new bout. We exported behavioral strings from BORIS to the BORIS tool Behatrix (http://www.boris.unito.it/pages/behatrix), in which we conducted a 10,000 permutation test to limit the behavioral transitions to those that occurred more frequently than chance (p < 0.05, transition occurring at least 1% of the time).

**Figure 2.**
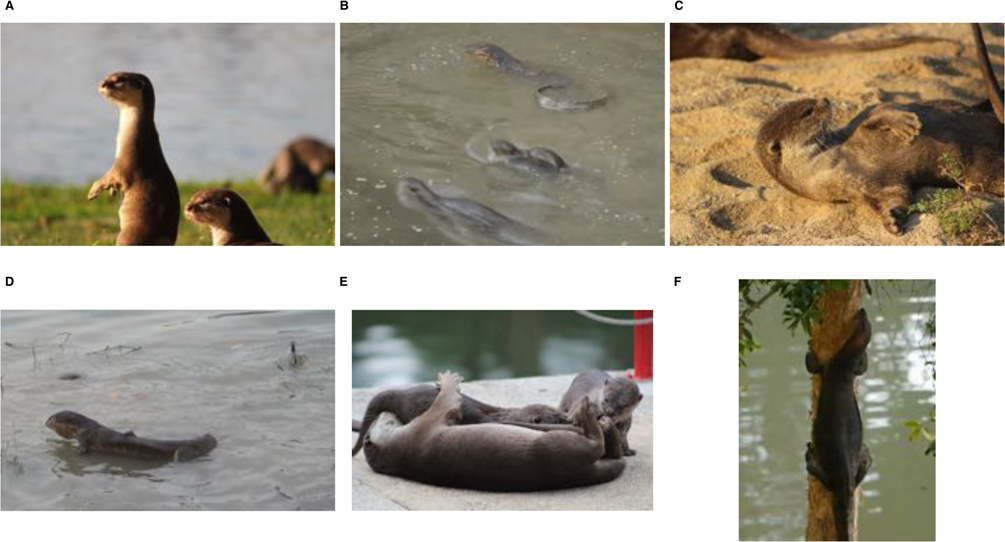
Notable otter and monitor behaviors. Otter vigilance (A), monitor lizard curling its tail (B), otter grooming (C), monitor with throat frill extended (D), otters playing (E), monitor climbing (F). See Table 1 for ethogram describing behaviors.

**Table 1.** Ethogram used in scoring otter and monitor behaviors.

| Behavior | Type | Description | Category |
| --- | --- | --- | --- |
| Fight | State | Monitor eng | aggression |
| Escape | Point | Monitor quic | locomotion |
| Climb | Point | Monitor clim | locomotion |
| Curl | State | Monitor curl | sound/sign |
| Bite | Point | Monitor bite | aggression |
| Static | State | Monitor is st | sound/sign |
| M_come | State | Monitor mov | locomotion |
| M_go | State | Monitor mov | locomotion |
| Whip | Point | Monitor whi | aggression |
| Whip hit | Point | Monitor whi | aggression |
| Frill | Point | Monitor frills | sound/sign |
| Mouth | State | Monitor oper | sound/sign |
| M_lunge | Poin | Monitor acce | aggression |
| M_eat | State | Monitor eati | sound/sign |
| O_bite | Point | Otter bites r | aggression |
| O_come | State | Otter moves | locomotion |
| O_eat | State | Otter in act c | sound/sign |
| O_go | State | Otter moves | locomotion |
| O_lunge | Point | Otter quickly | aggression |
| Play | State | Otter playfig | sound/sign |
| Groom | State | Otters groon | sound/sign |
| Sit | State | Otter sits stil | sound/sign |
| Touch | Point | Otter contac | aggression |
| Vigilance | State | Otter strikes | sound/sign |
| Retreat | Point | Otter reacts | locomotion |
| Float | State | Otter is stati | sound/sign |
| Dive | Point | Otter dives u | sound/sign |

For each video and bout we also recorded: presence of pups, number of pups if present, number of adults, number of monitors, age of pups, whether the events happened on land or water, whether there was an aggressive interaction, location of interaction, time of day, relative size of monitor to otter, minimum distance between otter and monitor, and general activity of otters (Table 2). We then constructed generalized linear mixed models (GLMMs), where we examined each bout as unique (n=234), unless it occurred both on land and in water, for a total of 270 analyzable interactions. We considered general movement between land and water benign behaviors, meaning the bout would end and a new one begin, should this occur. We considered the presence or absence of otter aggression, per otter, per bout. We examined otter vigilance with linear mixed models (LMMs), where we treated the initiation of otter vigilance as a point event, and because bouts varied in length, we treated vigilance as a continuous rate, per otter, per bout-minute. For consistency, we analyzed vigilance using the same subset of potentially aggressive encounters, even though vigilance could potentially occur when otters and monitors were further than one body length apart.

**Table 2.** List of variables used in GLMMs and LMMs.

| **Variable** | **Description** | **Type of Effect** |
| --- | --- | --- |
| **Adults** | Number of adults in observation | Fixed |
| **Pups** | Number of pups in observation | Fixed |
| **Pup Age** | Age of pups in days since first holt emergence | Fixed |
| **Environment** | Whether interaction occurred on land or in water | Fixed |
| **Environment/Pups** | Cross interaction of environment and number of pups | Interaction |
| **Adults/Pups** | Cross interaction of number adults and number of pups | Interaction |
| **Bout ID** | Unique bout ID | Random |

We generated GLMMs and LMMs with functions glmer and lmer, depending on whether the dependent variable was binomial (otter aggression) or continuous (vigilance), using the R package (R Core Team 2021; https://www.R-project.org/) lme4 (Bates et al. 2020). Fixed and random effects are in Table 2. We visualized flow diagram scripts from BORIS using the R package “DiagrammeR” (Iannone 2020), and we used the Tidyverse package (Wickham & RStudio 2019) for data analysis and created figures with the ggplot2 and sjPlot packages. Because this is a descriptive study, we present multiple linear models and then discuss their common features.

## Results

Monitor lizards displayed aggressive behaviors in more videos than otters did. Monitors displayed aggression in 79% (50/63) of videos, but otters were generally not overtly aggressive, displaying aggressive behaviors in only 27% (17/63). We found a significant association between otter and monitor aggression among videos (Fishers exact P = 0.013). In all videos where otters displayed aggression, monitors also displayed aggression (17/17); but otters were only aggressive in 34% of the videos where monitors were aggressive (17/50). Monitors displayed aggression in 72% of videos where otters did not (33/46), and otters never displayed aggression when monitors did not (0/13). Including only videos where otters and monitors were within a body length, monitors almost always displayed aggressive behaviors (33/34; 97%) while otters did only half the time (17/34).

To gain more insight into the causes of aggression, we examined the behavioral sequences within bouts. Permutation tests on behavioral sequences revealed that otter aggression typically led to monitor aggression, and not the other way around (Figures 3 & 4). For example, otters floated near, dived, then attacked monitor lizards (Figure 3, arrow a). Aggressive otter behaviors included biting or touching the head or tail of the monitor, which often responded by either curling its tail and frilling its neck, or by whipping its tail (Figure 3, arrows b, c & d). Although monitors often whipped their tails when attacked, they did not always hit; on land and in water (Figures 4A & 4B), monitors were about four times as likely to whip their tails after otters touched them as to successfully connect (22.2% vs 5.6%). Many aggressive behaviors by otters did not have any significant monitor precursors, but rather were starting points for a sequence of aggressive behaviors (e.g., Figure 4A, arrow a). In the water, otters often dived and emerged next to the lizard, which led to the lizard escaping away (Figure 4B, arrow d). Some behaviors frequently led to one another, such as lizards frilling and curling (Figure 4A, arrow b), or otters alternately displaying vigilance and playing (Figure 4A, arrow c). (For the purpose of this study we did not distinguish among play behaviors.)

**Figure 3.**
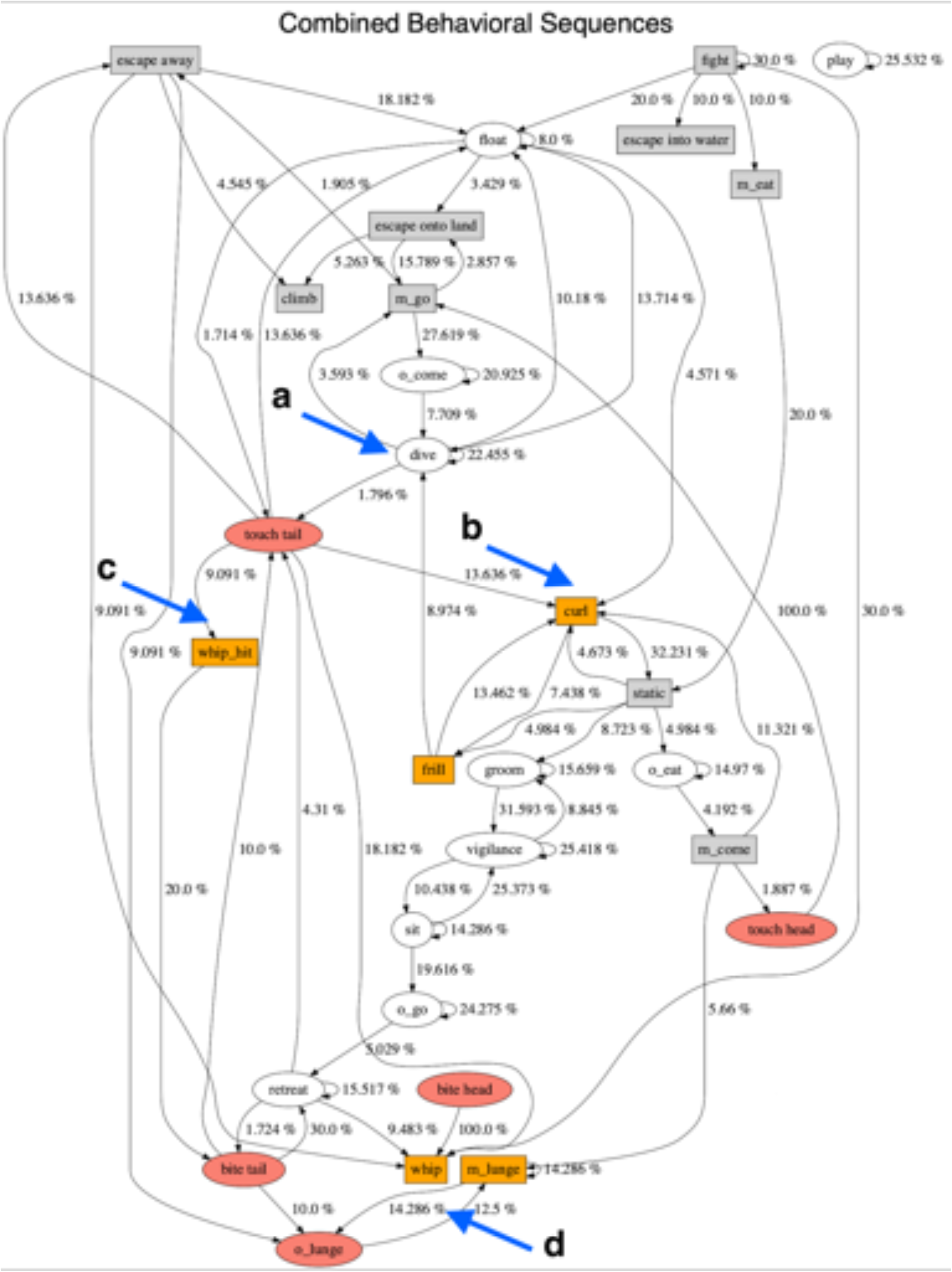
Behavioral sequences of all otter-monitor interactions. Ovals represent otter behaviors; rectangles represent monitor behaviors; colored symbols indicate aggression. See Table 1 for ethogram. Numbers describe transitions between behaviors; only transitions that occur more frequently than chance are shown. Letters indicate behaviors that precede aggression (see text).

**Figure 4.**
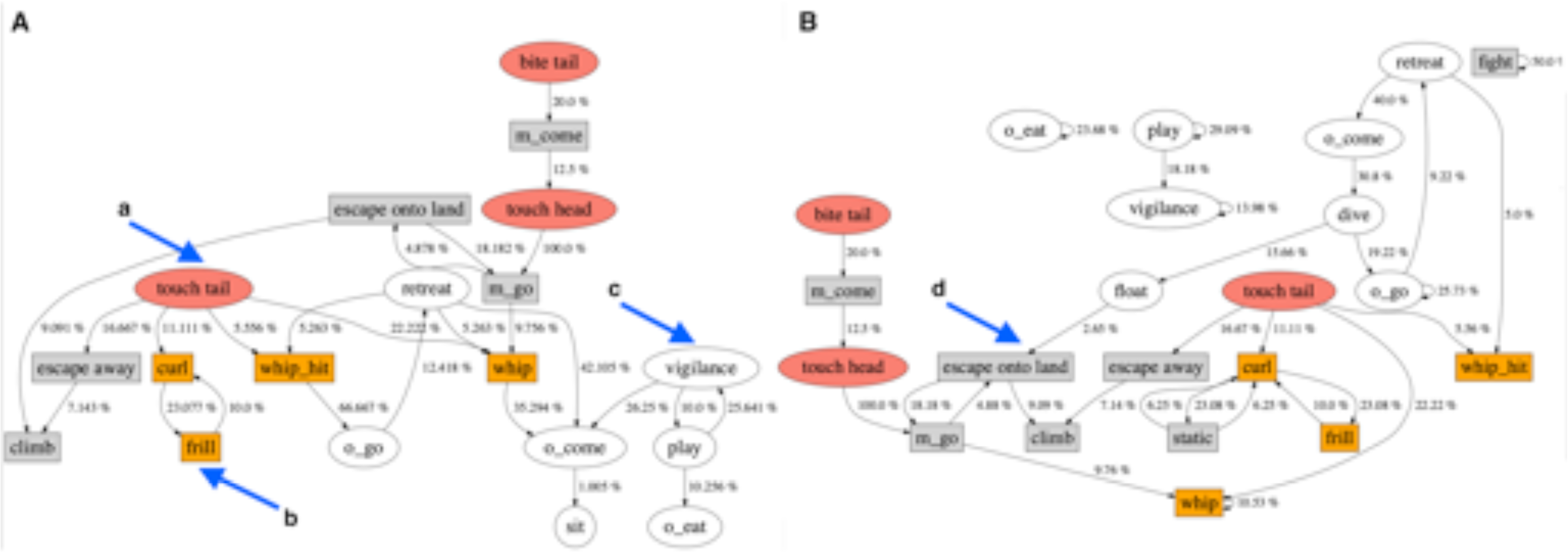
Behavioral sequences of otter-monitor interactions in water (A) and on land (B). Ovals represent otter behaviors; rectangles represent monitor behaviors; colored symbols indicate aggression. See Table 1 for ethogram. Numbers describe transitions between behaviors; only transitions that occur more frequently than chance are shown. Letters indicate behaviors that precede aggression or notable behavioral sequences (see text).

We examined whether the composition of otter groups within bouts influenced whether otters were aggressive. We found otters were more likely to display aggression towards monitor lizards in otter romps with pups than in romps without pups (Figure 5; Fisher’s Exact test p = 0.0018). We further examined the effects of group composition on the occurrence of otter aggression by fitting GLMMs to the number of pups, adults, pup age, and the environment (land or water) (Table 2). In all models, the number of pups in a group was a significant predictor of otter aggression (Table 3; Figure 6); groups with more pups were more likely to be aggressive. In the best model (Model 1; AIC = 59.1), the number of adults, the interaction between number of pups and adults, and pup age, were also significant predictors of otter aggression (Figure 6). However, in the simplest model, Model 2 (AIC = 60.6) only pups predicted otter aggression (Figure 7A). In Models 3 and 4 the number of adults also played a role in predicting otter aggression. In Model 3 (AIC = 62.3) the coefficient estimates for the number of pups and adults suggests a sigmoidal equation,

**Figure 5.**
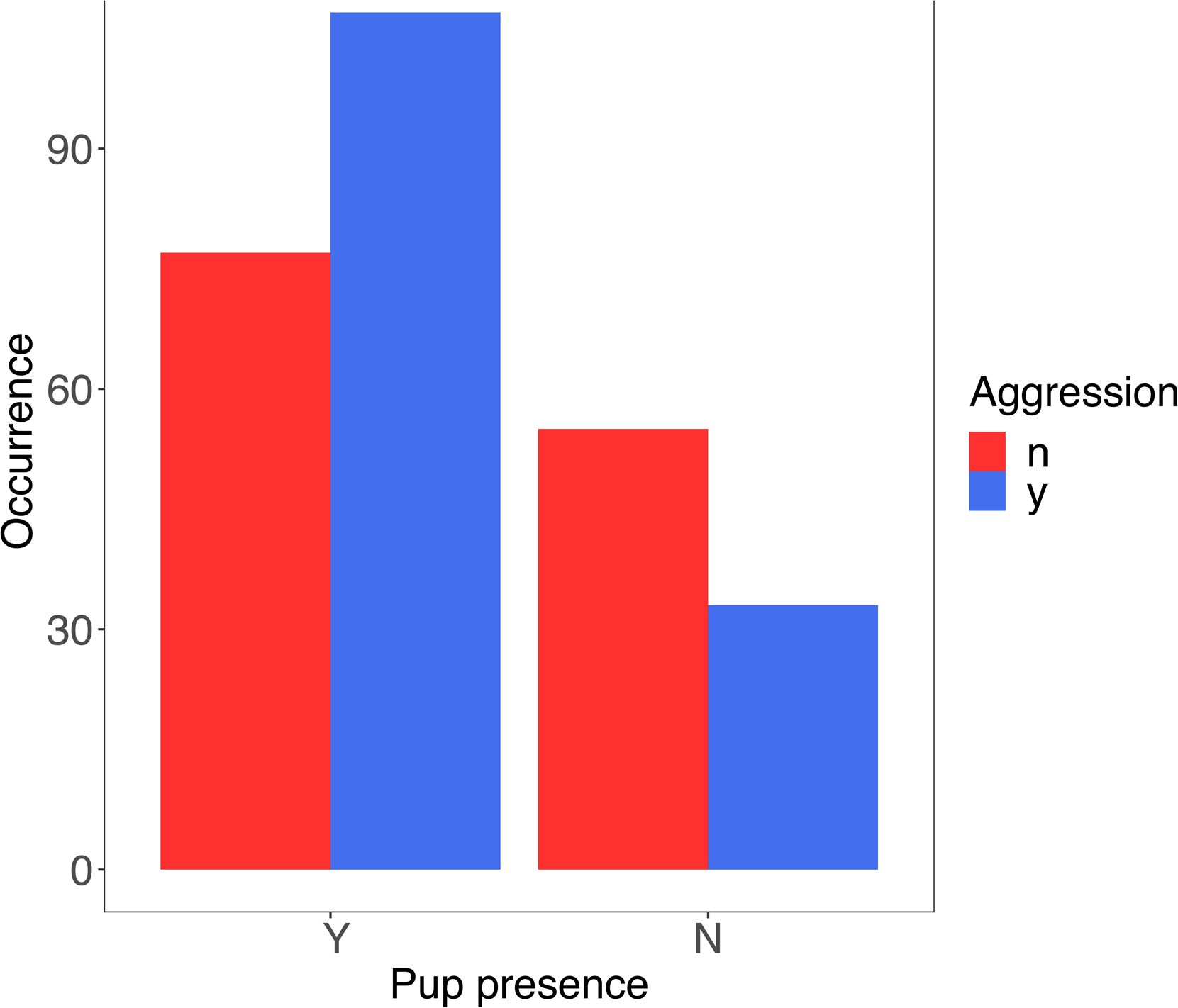
Occurrence of bouts with (y) and without (n) otter aggression in otter groups with (Y) and without (N) pups.

**Figure 6.**
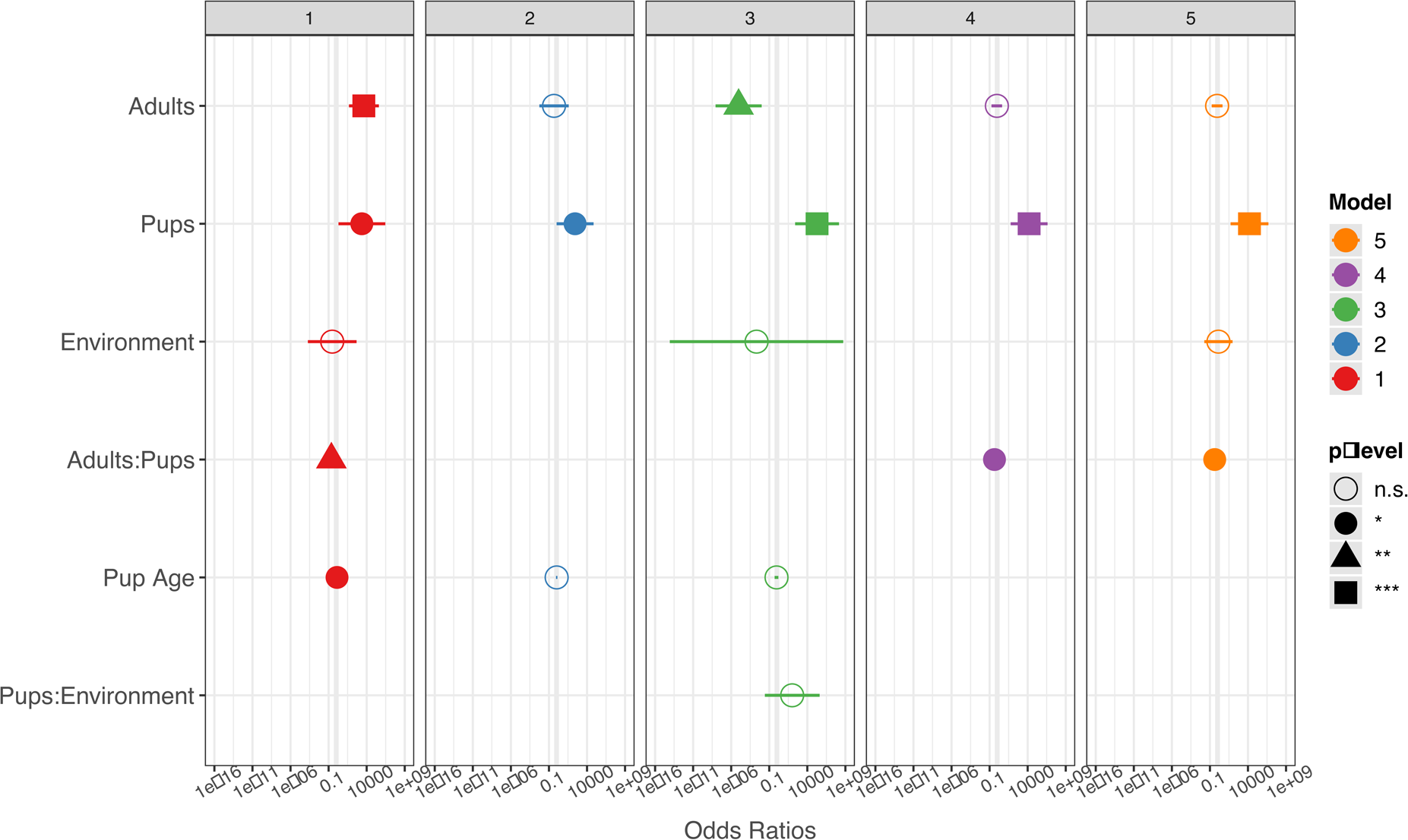
Coefficients for GLMM models 1-5 explaining occurrence of otter aggression.

**Figure 7.**
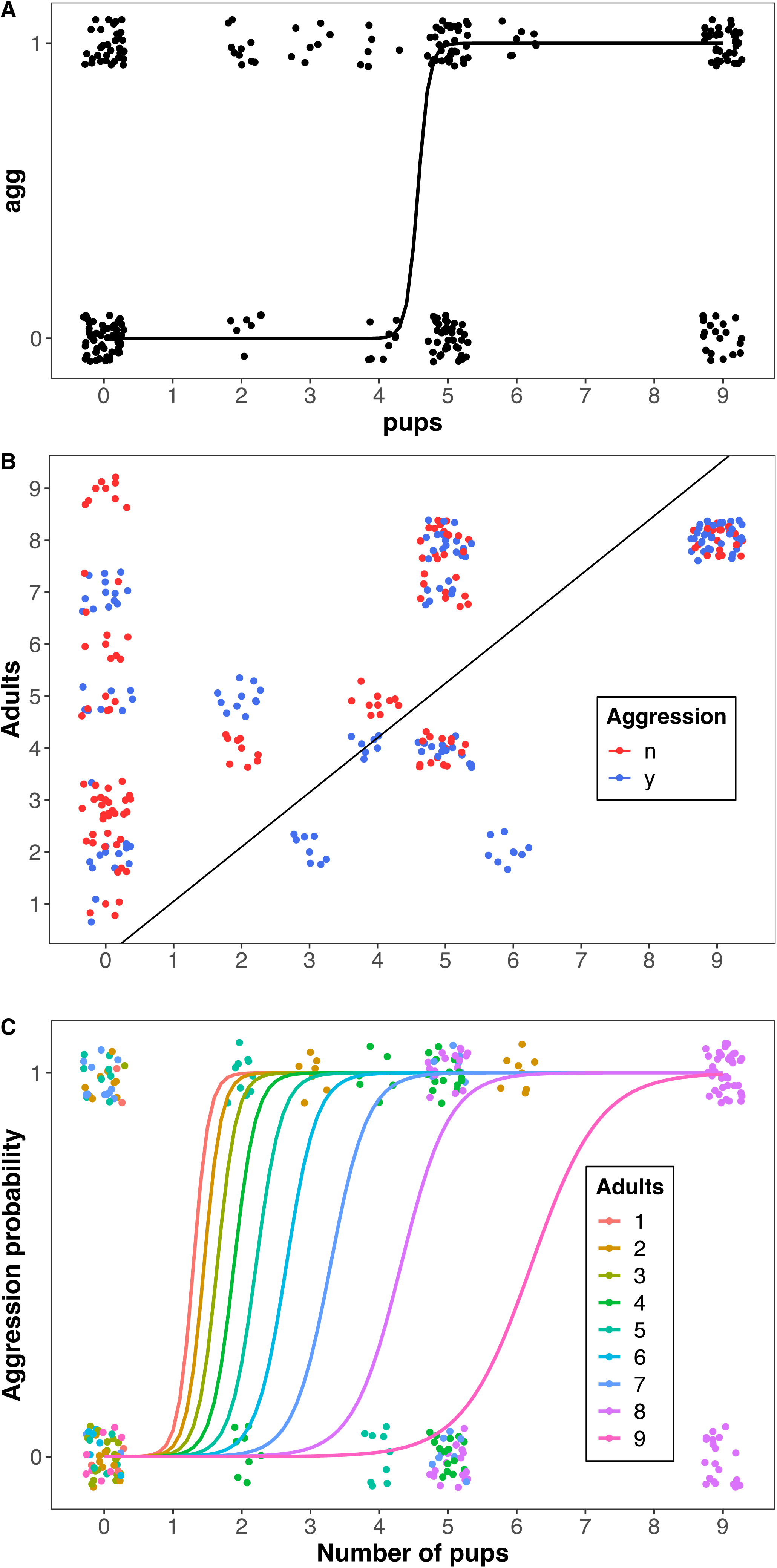
Models 2, 3, and 4 (panels A, B, C, respectively) predicting otter aggression on the number of pups in a group (X-axis; see Figure 6). In panels A and C, Y-axis represents probability of otter aggression; in panel B Y-axis represents the number of adults in a group, with a line of inflection points, adults = 1.05*pups, to the right of which otters are more likely to be aggressive (see text).

**Table 3.**
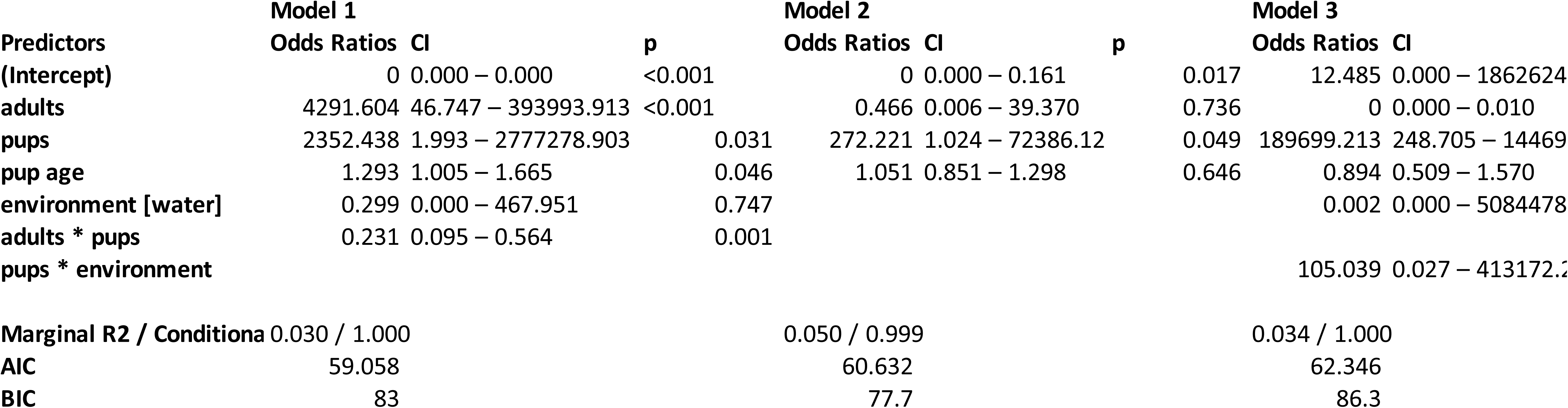
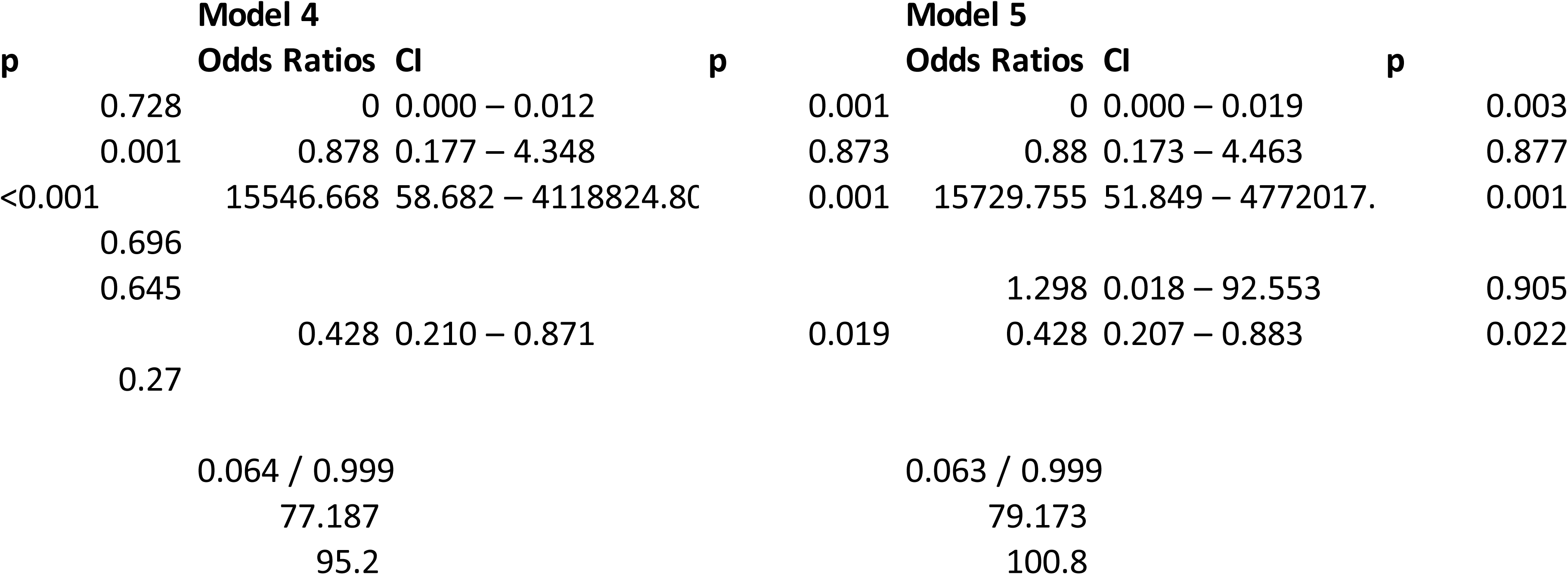
Generalized linear mixed models 1-5 predicting otter aggression towards water monitors.

Z = -11.58*Adults + 12.15*Pups

which implies a line of inflection points for the binomial GLMM with a slope,

adults = 1.05*pups

This result suggests that if there were more pups than adults in a group, adults were likely to display aggression towards monitor lizards (Figure 7B; Appendix 2), and therefore, groups with more adults required more pups before becoming aggressive. Model 4 includes the interaction between the number of pups and adults as a significant predictor of otter aggression (Figure 7C). Although Model 4 (AIC 77.2) has less support than Models 1, 2, and 3, the effect of adults is qualitatively similar to that in Model 3: more adults in a group shifted the inflection point for otter aggression (Figure 7C). Pseudo-R^2^ values (Nakagawa and Schielzeth 2013) for these models ranged from 0.031-0.064 (Table 3), but notice that low pseudo-R^2^ values are typical of logistic models (Hosmer et al. 2013).

### Vigilance

We examined what influenced otter vigilance by fitting the same fixed and random effects (Table 2) to LMMs explaining the rate of vigilance (Table 4). The first three vigilance models performed similarly (AIC = 650.0 to 653.3; Table 4); and some combination of pups, environment, and pup age were significant predictors in all four models. The number of pups in a group was a significant predictor in 3 out of the 4 models (Figure 8). There was a difference between vigilance rates on land and water (Figure 9A); adult otters displayed higher vigilance levels on land. Pup age was a significant predictor of otter vigilance in Model 4 (Figures 8, 9B). All four models have pseudo-R^2^ values above 0.3, indicating reasonable explanatory power, especially for an unmanipulated observational study (Table 4).

**Figure 8.**
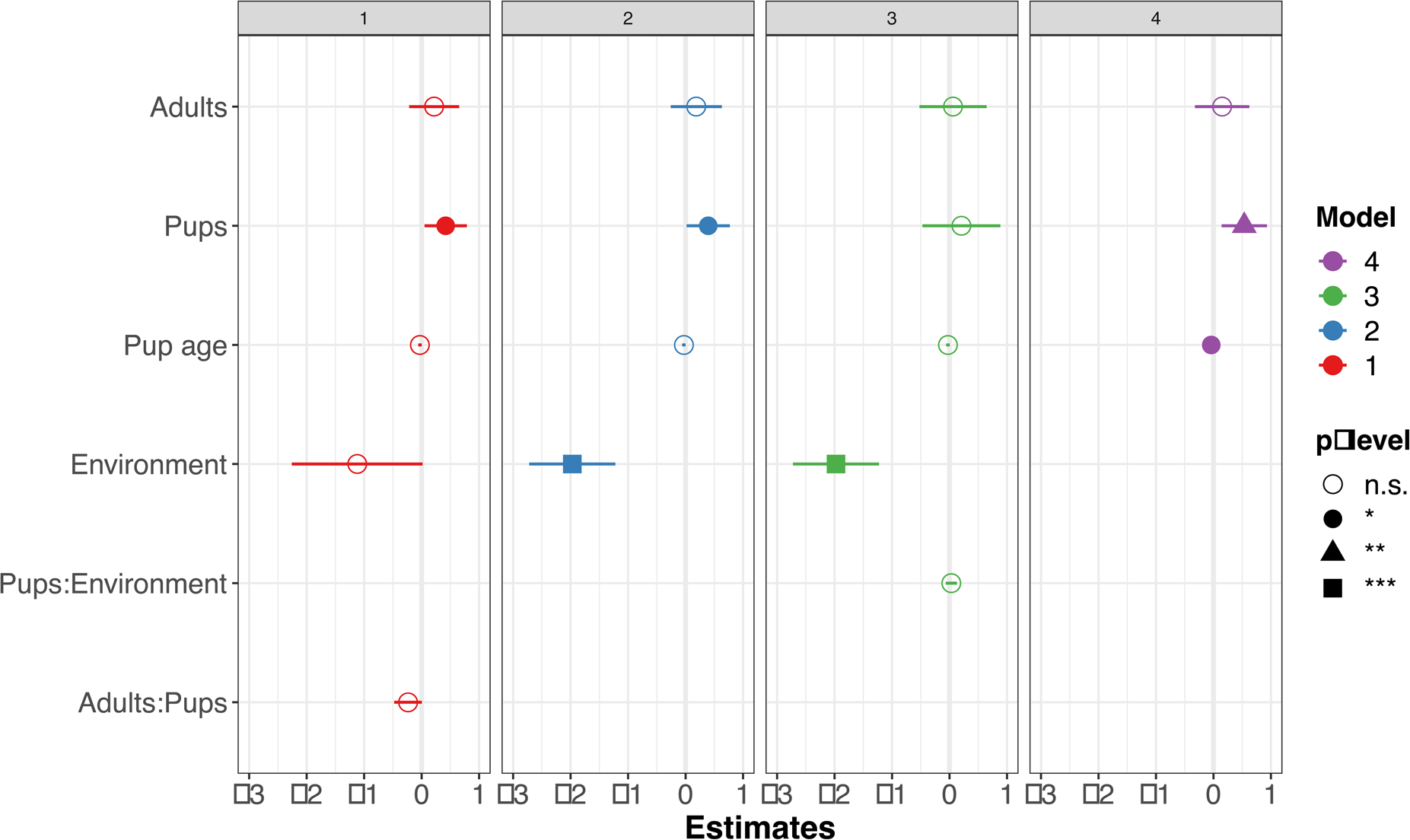
Coefficient estimates for LMMs, models 1-4, explaining the rate of vigilance, per otter, per minute.

**Figure 9.**
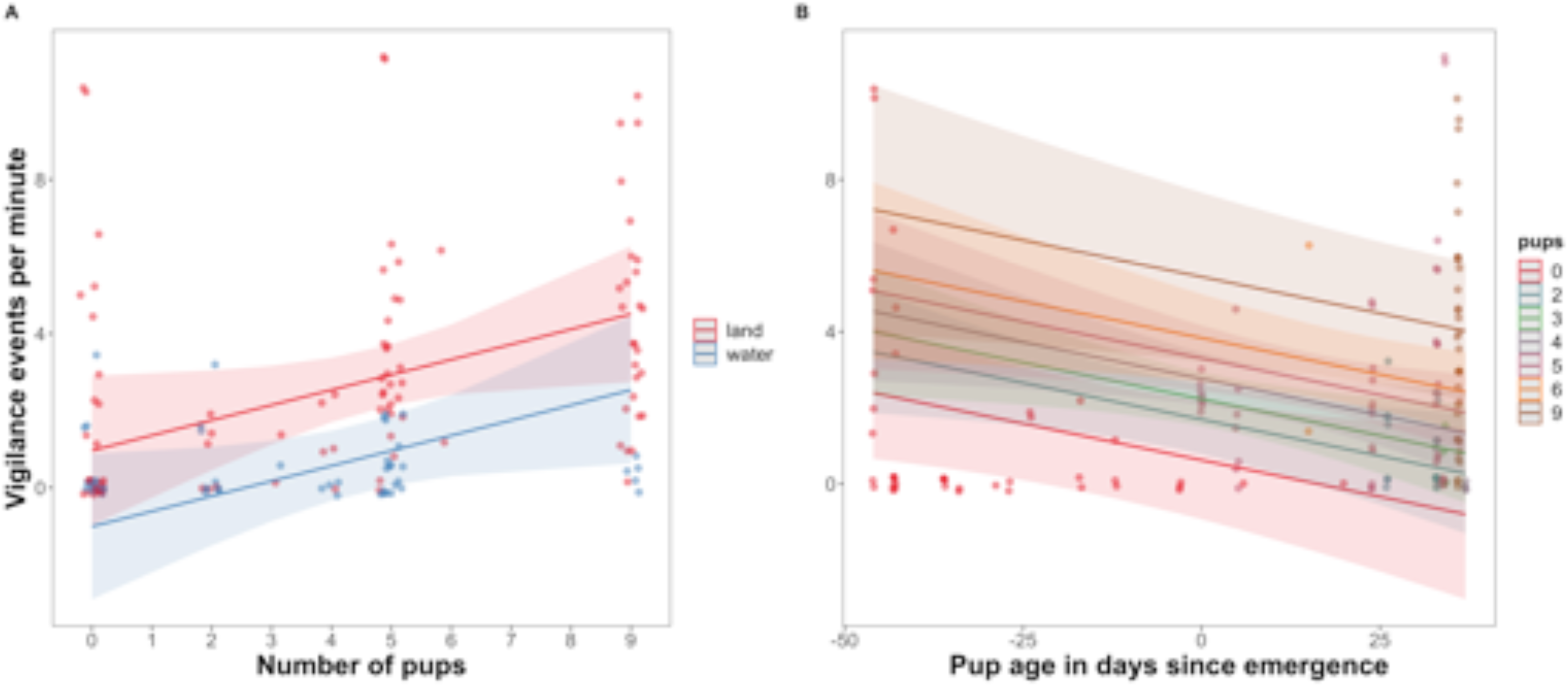
Models 2 and 4 (panels A and B, respectively) predicting rate of vigilance on the number of pups in a group (see Figure 8). Panel A includes environment (land or water); Panel B includes number of adults in a group (X-axis). Shaded areas are 95% confidence intervals.

**Table 4.**
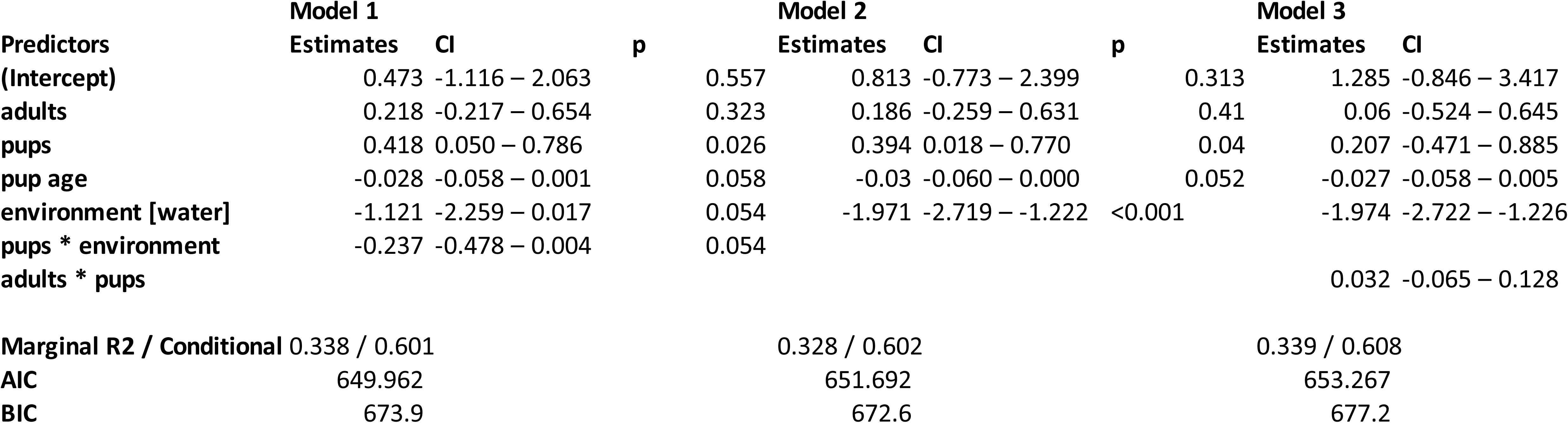
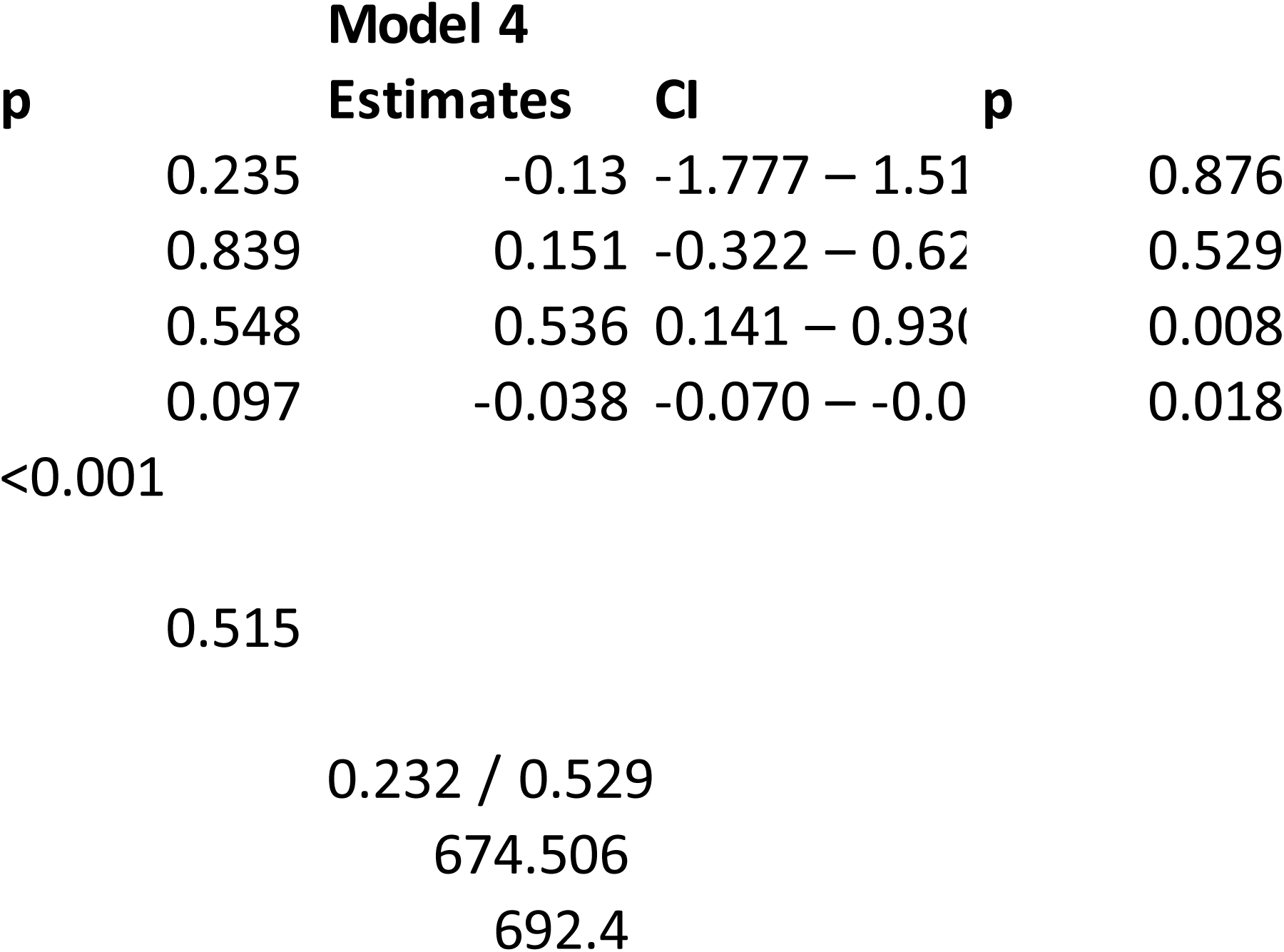
Linear mixed modelss 1-4 predicting rate of otter vigilance in the presence of water monitors.

## Discussion

In interactions between otters and monitor lizards, we found monitor lizards were aggressive or defensive far more often than are otters. Monitors adopted defensive postures, consisting primarily of a frilled neck and curled tail, even when otters were too far away to cause any physical harm, and monitors were especially aggressive or defensive when otters were within range of an attack (33/34 observations). The frequency of defensive monitor lizards suggests that otters are not responding to monitors, usually, but that monitors respond to the otters’ presence and approach. In general monitors react to otters as a threat.

Our behavioral sequence analysis supports this conclusion. Many of the monitors’ defensive and aggressive behaviors are preceded by an otter behavior, notably otters biting or touching monitors’ tails (Figures 3 & 4), and in water these behaviors are often preceded by diving or floating nearby (Figure 4). Monitors nearly always frill their necks and curl their tails when an otter is near them on land, even when the otter does not directly approach the lizard. Behavioral sequence analysis revealed no single cue from lizards that caused otters to react aggressively; nor was a cue obvious from our personal observations. Some behaviors led to predictable outcomes, e.g., if a monitor whipped its tail at an otter, then the otter retreated, but we found no behaviors that predicted why the otters first attacked the lizard.

Adult and subadult otters are fast and extremely agile swimmers, much more so than monitor lizards, even though monitors spend much of their day in water. Monitors frequently exit waterways and head onto land when otters are near; we observed this in 20 of the 63 scored videos, while a monitor escaped into the water only once. Once on land, monitors typically stay in place and adopt defensive postures. These behaviors suggest that monitors are less vulnerable to otters on land than in water, even when otters have a numerical advantage. Monitors can quickly scurry away on land, and their tail-whips are less affected by water, possibly increasing their effectiveness on land. Monitors on land are also likely able to better view the entire otter romp and avoid being attacked from below the water surface, which occurred fairly often. Perhaps most importantly, otters cannot submerge monitors on land and kill them, which was observed twice in interactions that occurred after the study period (e.g., Mitchell 2021). In one of the fatal interactions observed, a second, much larger lizard was in the water prior to the attack, and very slowly swam away as more and more otters attacked the smaller lizard. The larger lizard may have been just as vulnerable as the smaller lizard, despite its size, and took actions to leave the scene.

Monitors generally reacted the same way to otters, especially when otters were within one body length. The consistent defensive reaction of monitors suggests that the lizards do not primarily use pups as a cue. However, anecdotal evidence suggests monitors can distinguish pups from adults, including a monitor lizard attacking of a pup (Lee 2019) as well as one chasing a pup in one of the 63 videos in our study. There, a medium-large lizard chased a pup into the holt, with a few adult otters nearby. The pup had emerged from the holt about one month earlier, suggesting it was around 10 weeks old, and it was able to outrun the lizard. The monitor then moved away with no aggressive reaction from the adult otters. Presumably the monitor’s actions were predatory, and it is not clear why the adult otters did not react in this case.

### Otter aggression

Overall, we found that otters were only aggressive in encounters where monitors were also aggressive. However, that included situations where the otters instigated an attack, and the monitors responded, i.e., otters are not responding to monitor aggression *per se*. This observation supports the notion that monitors are reacting to the presence of the otters, rather than otters reacting to the behaviors of the lizards. Our sequential analysis did not give a clear indication of a “trigger” for otter aggression, other than a monitor approaching the otter head on, which was a rare occurrence (3.8% of transitions). Although otter aggression towards monitor lizards can be lethal, in general otters are not especially aggressive, displaying aggression half the time an attack was possible.

In all the models we examined, the number of otter pups in a group contributed to otter aggression towards monitor lizards. Otters are less likely to be aggressive in the absence of pups, and otter aggression is more likely as the number of pups increases, especially if there are more than five pups. When more pups present, there is a greater chance that at least one of them will encounter a threat. Otter pups are notoriously curious and active; they show little hesitancy to approach or even touch a monitor lizard. We generally observed at least one adult otter in the immediate vicinity of pups, but there were times when it seemed as if the adults could not oversee all the pups at once. One pup might wander near a monitor, and an adult would then attack the lizard; this was the case in one of the fatal incidents we observed (below).

One observation that occurred after our study period was especially illustrative. A small monitor was attacked and partly eaten by the romp of 20 otters at Ulu Pandan (this was the Jurong Lake Gardens romp; they displaced the original Pandan romp in early 2020). Here, a pup approached and contacted the lizard, which whipped its tail, leading to the ensuing melee. Pups in the romp were very young, having only emerged from the holt about one week prior. After a series of tail-whips kept the adult otters at bay, the lizard maintained its defensive stance as the otters appeared to regroup nearby on the bank. Notably, one adult otter began carrying the pups away from the lizard, towards the holt, with the pups scurrying back down in between drop-offs. After a minute of attempting to shift pups and milling around, one adult charged the monitor and the rest followed suit. The otters flipped the monitor onto its back and dragged it into the river, where they continued to attack it for over 30 minutes. They later appeared to begin eating the floating corpse. This occurrence supports the idea that the lizards are vulnerable to large romps of otters, even romps with young pups. It builds on Goldthorpe et al.’s (2010) observation, which speculated on but did not confirm the death of the lizard and subsequent consumption by the otter.

Adult numbers play a role as well. In two of the five models predicting otter aggression, the number of adults had a significant effect, and in three models the interaction between the number of adults and pups had a significant effect (Table 3). The combined effects of adults and pups were qualitatively similar across models (Figure 6, Appendix 2): if few adults are present, having only a few pups in a group can lead to otter aggression; with more adults in a group, more pups need to be present to lead to aggression. Small groups of adults can be aggressive. Even lone otters sometimes attack monitors and even larger species, such as estuarine crocodiles (Goldthorpe et al. 2010). One explanation for the relationship between adults and aggression is that larger numbers of adults are a natural deterrent to monitor attacks. Consequently, in groups with more adults, otters may be less prone to react to monitors as threats. Another possibility is that, with larger numbers of adults, there are fewer wayward pups. Larger groups may have more babysitters, effectively. We observed groups with wide ranges of adult and pup numbers, including several adults with few pups, so the pattern is unlikely to be an artefact of group size, per se.

Group composition itself could play a role, and which adults are present may determine whether otters are aggressive or not. For example, if certain individuals such as the breeding pair initiate attacks on monitor lizards, then additional adults may not increase the chance of aggressive responses to monitors and may even dilute the breeding pair’s ability to respond to monitors. We cannot tease apart the effects of individual otters without being able to identify individuals, which we could not do in this study. Some evidence suggests that different otters have different tendencies to attack, however. In one bout, a large monitor lizard chased an otter pup under a human-made structure while several adult otters watched, but none reacted aggressively. Adult offspring may have been waiting for another otter to respond to monitors or may not have known what to do.

Pup age may also play a role. We found pup age was positively related to otter aggression in the best model (Figure 6) but negatively related to otter vigilance (Figure 8). Presumably at some point pups become independent enough that adults do not need to watch over them, but it is hard to reconcile these contrasting patterns in aggression vs vigilance. Pup age, like other factors in this observational study, warrants further investigation.

### Vigilance

In none of our models did the number of adults in a group have a significant effect on vigilance rates. This differs from many other studies of vigilance, where group size often decreases vigilance rates (see Quenette 1990, Fernández-Juricic E 2012). However we found that vigilance rates generally increase as the number of pups in a group increase, which supports the hypothesis that adult otters increase their vigilance rates to compensate for unequal pup contribution. The effect was significant in 3 out of our 4 models. Pups are curious and frequently venture near lizards; adults likely need to keep a better lookout. Vigilance is typically interspersed with other stationary riverbank behaviors, such as grooming and playing, but we found it did not by itself lead to aggression (Figures 3 & 4). Pup age was a significant factor towards increased vigilance levels in one model (model 4, Figures 3 & 4), but was either insignificant or marginal for the other three models. The duration of this study may not have been long enough to measure a change in adult behavior. At some point pups grow up, and presumably adults stop compensating for them then.

Vigilance rates were also higher when the otters were on land than in water. There could be several reasons for increased vigilance rates on land. One is that visibility is less obstructed on land. Otters often go onto banks to groom, eat, or visit spraint sites, and the group remains more stationary than in water, allowing otters to better “keep watch” over each other. But predators can spot otters, too, and otters on land may be more vulnerable to predators than swimming otters. The increased risk may compel otters to be more vigilant on land. On land, they are likely more vulnerable not just to large monitors, but also to packs of dogs and even crocodiles (Clements 2019), plus, of course, humans. Swimming otters’ agility seems to explain why lizards are reluctant to be in the water when otters are around; there are few animals in Singapore’s waterways that otters cannot outswim. The difference between land and water was probably not an artefact of our criteria for vigilance; we defined vigilance such that it could be observed on land and water.

Our study was not aimed at water monitor lizards, *per se*, but so little research exists on wild monitor behavior that it provided some insights. Otter watchers and other scientists had hunches that monitor lizards sometimes avoid smooth-coated otters, and our analyses bear this out: monitors respond defensively to otters. We observed only one incident of what was presumably a failed attempt by a monitor lizard to prey on an otter pup. Whether monitors regularly prey on otter pups is something we cannot ascertain from this study. Adult otters may respond aggressively to monitors, not specifically because they are a threat to pups, but because the monitors are large and live in the same areas. Anecdotally, groups of smooth-coated otters engage aggressively with larger species, such as dogs (e.g., Chua 2018), crocodiles (e.g., Choudhary 2019), and occasionally even tigers (Narasimhamurthy 2021), and romps of giant otters engage with caimans (Ribas et al. 2012) and jaguars (Leuchtenberger et al. 2016). Monitor lizards might simply do their best to avoid romps of angry otters.

Our findings derive from a study of a single romp of otters living on a highly modified river in Singapore, the Ulu Pandan River, where otters and monitors are both common. The extent to which our conclusions are generalizable to other families within Singapore, and to locations outside of Singapore, is not clear. Collecting animal behavior data is very time and labor intensive by nature, and here we were able to glean information from videos collected by local otter watchers. By crowdsourcing video recording, citizen science has the potential to be an extremely powerful tool in animal behavior studies due to the “many eyes” effect: if more people are recording animal behaviors, we can collect a more complete record of what animals do. In this manner, this analysis of data gleaned from the internet can be viewed as a form of “next-gen” natural history (Tosa et al. 2021). But this approach has limitations. The current study gleaned information from videos collected in an *ad lib* manner, albeit with impressive regularity. (The only gap in otherwise daily video recordings of wildlife was about three weeks during the COVID-19 lockdown). Biases in *ad lib* data collection can skew data towards rare, conspicuous behaviors. However, our videos were taken at about the same time and place every day, which should reduce bias towards any particular set of behaviors. Further, we limited which videos we included in the analysis to those with otters and monitors. We make no claims about the overall frequency of otter-monitor interactions (other than that they are surprisingly frequent) and limit our analysis to what happens during otter-monitor interactions. We hope the regular, frequent, and relatively unbiased collection of videos, combined with our filtering the videos to a particular narrow topic, reduced unintentional biases.

## Conclusions

While we did not discover a specific behavioral “trigger” that otters use as a cue to attack monitor lizards, we did find several factors that affect the likelihood of aggression towards a monitor. Otters were only aggressive in 50% of close-up interactions. Monitors, conversely, displayed aggressive or defensive behaviors in almost all such encounters. Otters are faster and more agile, especially in water, and combined with their group behaviors, can pose a real threat to monitors. Monitors seem to be content to scavenge what the otters leave behind and otherwise avoid them entirely if possible. The presence of young pups increases the chance that otters act aggressively to monitors and increases the rate of vigilance within the otter group. The increased vigilance rates could then lead to otters being more aware of monitors’ presence, with a greater chance of aggression resulting. To what extent the growing population of large otter romps in Singapore’s very urban environment contributes to this is something we cannot ascertain from this study.

## Appendix legends

**Appendix 1.**
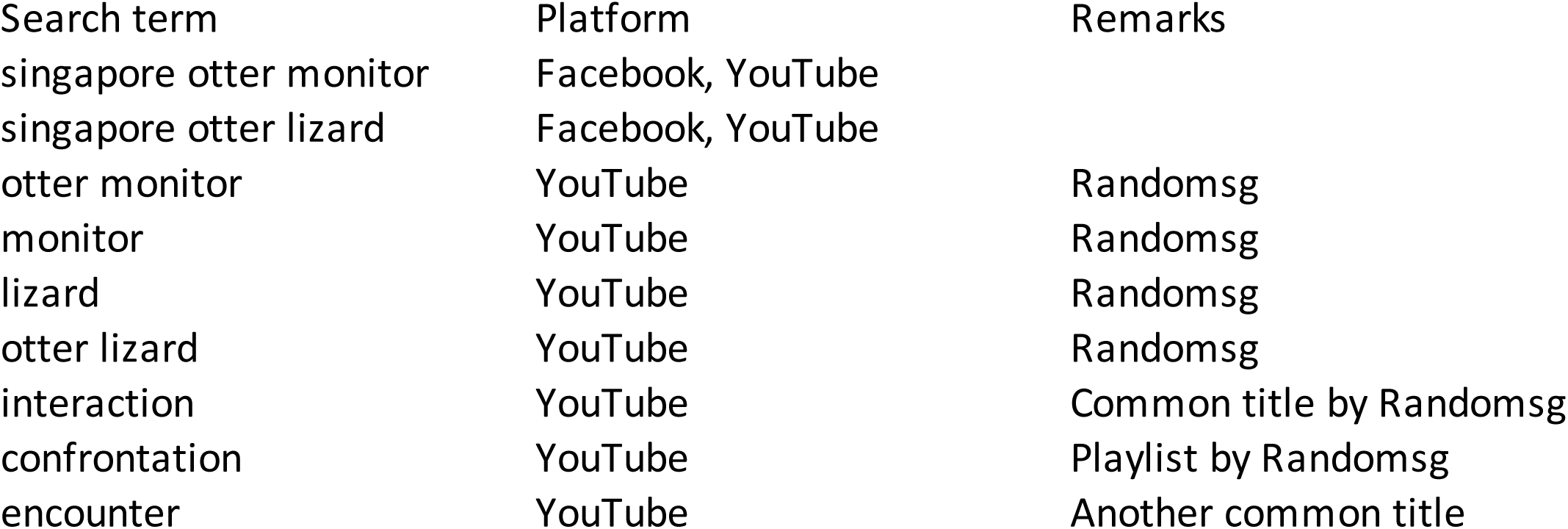
List of search terms used on social media to find otter-monitor videos.

**Appendix 2.**
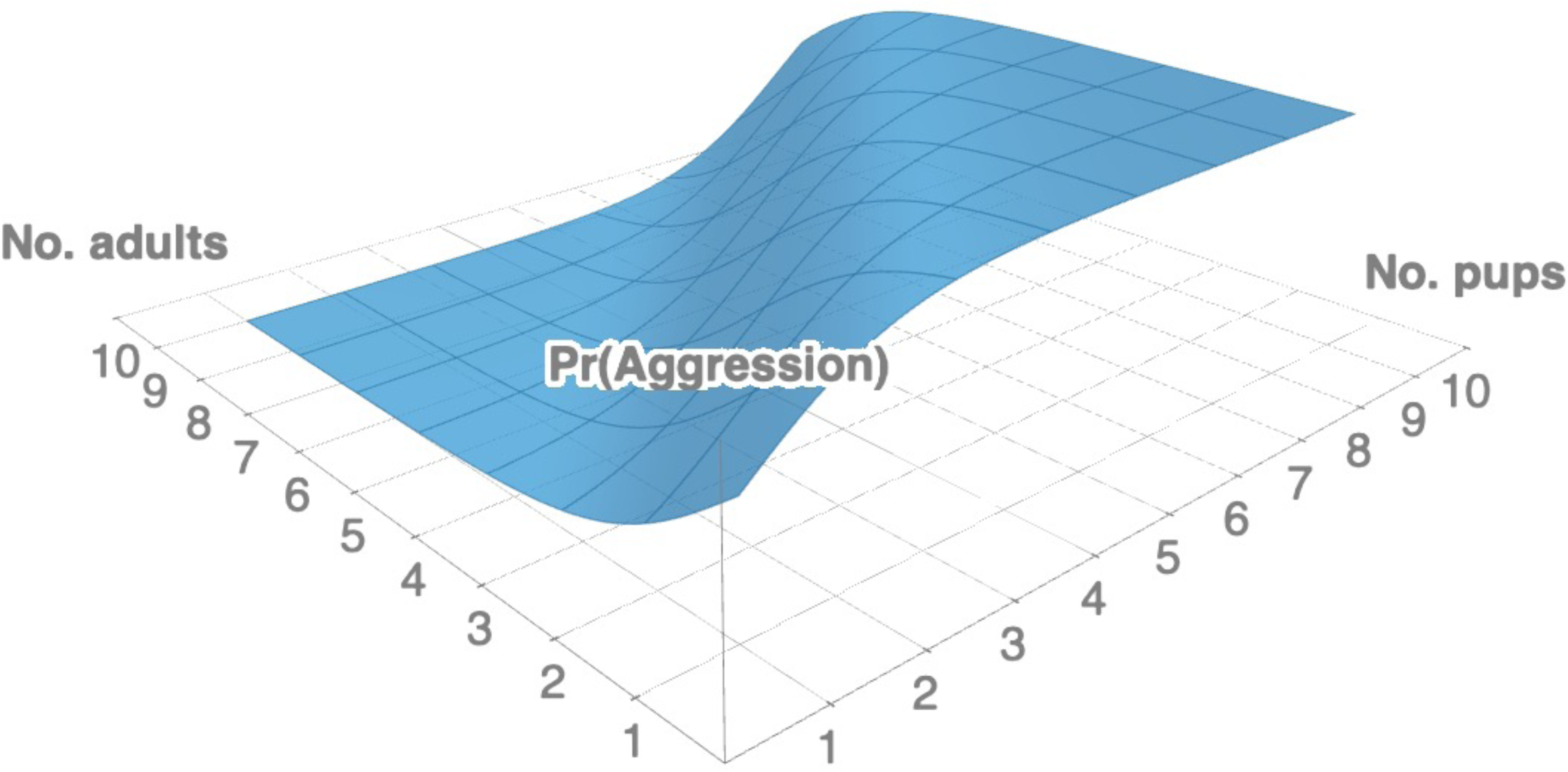
Three-dimensional depiction of the function described by Model 3 (Figure 7) predicting otter aggression towards monitors as a function of pups and adults. See text for sigmoidal equation.

## Data accessibility

Raw data, BORIS files, and R code archived at FigShare, https://figshare.com/articles/dataset/Bungum_Johns_Otter-MonitorLizard-Interaction-Data/20188241, https://doi.org/10.6084/m9.figshare.20188241

## Acknowledgments

We would like to thank the otter-watching communities of Singapore for their helpful comments and conversations, especially Alicia Ellen Brierley, Max Khoo, Alex Mitchell, Sivasothi N, “Uncle George” Wong, Sabrina Wong, and the YouTube channel RandomSG. We would also like to thank Bree Putman for her helpful input on an early draft of this manuscript. This study was conducted in partial fulfilment of the Yale-NUS College Capstone requirements for Haaken Bungum. The work was supported in part by the Ministry of Education through the Yale-NUS College start-up grant R-607-265-226-121, by Yale-NUS College Centre for International and Professional Experience (CIPE), and by NParks permits NP/RP20-073 and NP/RP20-075.

## References

Altmann, Jeanne. 1974. “Observational Study of Behavior: Sampling Methods.” Behaviour 49 (3/4): 227–67. http://www.jstor.org/stable/4533591.

Azevedo, Olga, Ana Correia, Ana Magalhaes, and Liliana de Sousa. 2015. “How Do Common Otters (Lutra Lutra, Linnaeus 1758) Interact? Behavioral Study on a Pair of Otters in Captivity.” Animal Behavior and Cognition 2 (2): 124–31. doi:10.12966/abc.05.02.2015.

Bates, Douglas, Martin Maechler, Ben Bolker [aut, cre, Steven Walker, Rune Haubo Bojesen Christensen, Henrik Singmann, et al. 2020. “Lme4: Linear Mixed-Effects Models Using ‘Eigen’ and S4”. https://CRAN.R-project.org/package=lme4.

Beauchamp, Guy. 2001. “Should Vigilance Always Decrease with Group Size?” Behavioral Ecology and Sociobiology 51 (1): 47–52. doi:10.1007/s002650100413.

Beauchamp, Guy. 2008. “What Is the Magnitude of the Group-Size Effect on Vigilance?” Behavioral Ecology 19 (6): 1361–8. doi:10.1093/beheco/arn096.

Belanger, Michael, Clough Nicole, Nesime Askin, Tan Luke, and Carin Wittnich. 2011. “A Review of Violent or Fatal Otter Attacks.” IUCN Otter Specialist Group Bulletin 28 (January).

Bertram, Brian. 1980. “Vigilance and Group Size in Ostriches.” Animal Behaviour 28 (1): 278–86. doi:10.1016/S0003-3472(80)80030-3.

Boydston, Erin E, and Eric S Abelson. 2018. “Canid Vs. Canid: Insights into Coyote-Dog Encounters from Social Media.” Human-Wildlife Interactions 12 (2): 233–42.

Breck, Stewart W., Sharon A. Poessel, Peter Mahoney, and Julie K. Young. 2019. “The Intrepid Urban Coyote: A Comparison of Bold and Exploratory Behavior in Coyotes from Urban and Rural Environments.” Scientific Reports 9 (1):2104. doi:10.1038/s41598-019-38543-5.

Burnham, Kenneth P., David R. Anderson, and Kathryn P. Huyvaert. 2011. “AIC Model Selection and Multimodel Inference in Behavioral Ecology: Some Back-ground, Observations, and Comparisons.” Behavioral Ecology and Sociobiology 65 (1): 23–35. doi:10.1007/s00265-010-1029-6.

Bungum, Haaken Zhong, Mei-Mei Tan Heng Ye, Atul Borker, Chia Da Hsu, Philip Johns. 2022. Observations of multiple reproductive females in groups of smooth-coated otters. Ethology 128: 285–291. DOI: 10.1111/eth.13263

Chua, Peter. 15 June 2018. “Otters vs Dogs none were hurt in the making of this video.” Facebook: Nature Society (Singapore). Accessed 15 March 2022. https://www.facebook.com/groups/naturesocietysingapore/posts/10156291154698213/

Choudhary, Ashit. 12 February 2019. “Size does not matter. Otters are known for their notorious behaviour of how they challenge their opponents”. Facebook: IUCN SSC Otter Specialist Group. Accessed 15 March 2022. https://www.facebook.com/OtterSpecialistGroup/posts/2097860866961395

Chung, Lyn-Yi. 2017. “5-Year-Old French Girl Bitten by Otter at Gardens by the Bay - CNA.” Channel NewsAsia. https://www.channelnewsasia.com/news/singapore/5-year-old-french-girl-bitten-by-otter-at-gardens-by-the-bay-9820916.

Clements, Claire. 2019. “Dogs Vs. Otters: Who Will Win? Wild City: Singapore Saturday at 9pm BBC America.” BBC America. https://www.youtube.com/watch?v=qcZlKCjsWkQ.

Cooper, C. 2016. Citizen Science: How Ordinary People are Changing the Face of Discovery. The Overlook Press, Peter Mayer Publishers, Inc. New York, NY, USA.

DeLisle, Harold. 2007. “Observations on Varanus S. Salvator in North Sulawesi.” Biawak 1 (2): 59–66.

Elgar, Mark A. 1989. “Predator Vigilance and Group Size in Mammals and Birds: A Critical Review of the Empirical Evidence.” Biological Reviews 64 (1): 13–33. doi:https://doi.org/10.1111/j.1469-185X.1989.tb00636.x.

Fernández-Juricic E. 2012. Sensory basis of vigilance behavior in birds: Synthesis and future prospects. Behavioural Processes. 89(2):143–152. doi:10.1016/j.beproc.2011.10.006.

Friard, Olivier, and Marco Gamba. 2016. “BORIS: A Free, Versatile Open-Source Event-Logging Software for Video/Audio Coding and Live Observations.” Methods in Ecology and Evolution 7 (11): 1325–30. doi:10.1111/2041210X.12584.

Goldthorpe, Gareth, Chris Shepherd, Hogg Stephen, and Boyd Leupen. 2010. “Pre-dation of Water Monitor Lizard (Varanus Salvator) by Smooth-Coated Ot-ter (Lutrogale Perspicillata) in Peninsular Malaysia.” IUCN Otter Specialist Group Bulletin 27 (January).

Hosmer, David W, Stanley Lemeshow, Rodney X Sturdivant. 2013. Applied Logistic Regression, 3^rd^ ed. Wiley Series in Probability and Statistics. John Wiley & Sons, Hoboken, New Jersey, USA.

Hwang, Yeen Ten, and Serge Larivière. 2005. “Lutrogale Perspicillata.” Mammalian Species 2005 (786): 1–4. doi:10.1644/786.1.

Iannone, Richard. 2020. “DiagrammeR: Graph/Network Visualization.” https://CRAN.R-project.org/package=DiagrammeR.

Kamjing, Anucha, Dusit Ngoprasert, Robert Steinmetz, Wanlop Chutipong, Tommaso Savini, and George A. Gale. 2017. “Determinants of Smooth-Coated Otter Occupancy in a Rapidly Urbanizing Coastal Landscape in Southeast Asia.” Mammalian Biology 87 (November): 168–75. doi:10.1016/j.mambio.2017.08.006.

Khoo, M. D. Y., and B. P. Y.-H. Lee. 2020. “The Urban Smooth-coated Otters Lutrogale Perspicillata of Singapore: A Review of the Reasons for Success.” International Zoo Yearbook, July. doi:10.1111/izy.12262.

Khoo, Max, and N Sivasothi. 2018a. “Observations of the Variation in Group Structure of Two Urban Smooth-Coated Otter Lutrogale Perspicillata Groups in the Central Watershed of Singapore.” IUCN/SCC Otter Specialist Group Bulletin 35 (December): 148–54.

Khoo, Max, and N Sivasothi. 2018b. “Population Structure, Distribution, and Habitat Use of Smooth-Coated Otters Lutrogale Perspicillata in Singapore.” IUCN/SCC Otter Spe- cialist Group Bulletin 35 (December): 171–82.

Krueger, Konstanze, Laureen Esch, and Richard Byrne. 2019. “Animal Behaviour in a Human World: A Crowdsourcing Study on Horses That Open Door and Gate Mechanisms.” PLoS ONE 14 (6). doi:10.1371/journal.pone.0218954.

Lay, Belmont. 2021. “Otters Crossing Busy Orchard Road Outside Plaza Singapura Almost Hit by Taxi That Braked in Time.” News. Mothership.sg. https://mothership.sg/2021/01/otters-cross-orchard-road/.

Lee, Phyllis. 2016. “Sentosa Cove Residents Put up Fences to Fend Off Otters Preying on Fish in Homes.” Text. The Straits Times. https://www.straitstimes.com/singapore/otters-prey-on-fish-at-sentosa-cove-homes-desperate-residents-put-up-fences-motion-sensor.

Lee, Van Hien. 2019. “Malayan Water Monitor Attacked a Smooth Otter Pup – Bird Ecology Study Group.” Bird Ecology Study Group. https://besgroup.org/2019/07/18/malayan-water-monitor-attacked-a-smooth-otter-pup/.

Lee Wei Lin. 2020. “Former Actress Jazreel Low Doesn’t Want the Otters That Ate over 100 of Her Fishes to Be Culled; Says It’s ‘Not the Right Way Out’.” TODAYonline, May. https://www.8days.sg/sceneandheard/celebrities/former-actress-jazreel-low-doesn-t-want-the-otters-that-ate-over-12736180.

Leuchtenberger, Caroline, Samara Bezerra Almeida, Artur Andriolo, and Peter G. Crawshaw. 2016. “Jaguar Mobbing by Giant Otter Groups.” Acta Ethologica 19 (2): 143–46. doi:10.1007/s10211-016-0233-4.

Mitchell, Alex. 2021. Singapore Otter Project. YouTube. “Breakfast buffet -or perhaps an attack on a monitor?” 09 April 2021. Accessed 30 June 2022. https://www.facebook.com/alex.mitchell.79462815/videos/10159158842136678/

Nakagawa, Shinichi, and Holger Schielzeth. 2013. “A General and Simple Method for Obtaining R2 from Generalized Linear Mixed-Effects Models.” Methods in Ecology and Evolution 4 (2): 133–42. doi:https://doi.org/10.1111/j.2041-210x.2012.00261.x.

Narasimhamurthy, Harsha. 16 March 2021. “Tigers vs otter in water”. Twitter. Accessed 15 March 2022. https://twitter.com/HJunglebook/status/1371796984797958144

Nelson, Ximena J., and Natasha Fijn. 2013. “The Use of Visual Media as a Tool for Investigating Animal Behaviour.” Animal Behaviour 85 (3): 525–36. doi:10.1016/j.anbehav.2012.12.009.

Ng, Cherlynn. 2020. “Woman Injured After Being Bitten by Otter That ‘Charged’ Towards Her at Gardens by the Bay.” News. Stomp. https://stomp.straitstimes.com/singapore-seen/woman-injured-after-being-bitten-by-otter-that-charged-towards-her-at-gardens-by-the.

Quenette, Pierre-Yves. 1990. “Functions of Vigilance Behavior in Mammals: A Review.” Acta Oecologica 11 (January): 801–18.

R Core Team. 2021. R: A language and environment for statistical computing. R Foundation for Statistical Computing, Vienna, Austria. https://www.R-project.org/

Ribas, Carolina, Gabriel Damasceno, William Magnusson, Caroline Leucht-enberger, and Guilherme Mourão. 2012. “Giant Otters Feeding on Caiman: Evidence for an Expanded Trophic Niche of Recovering Popu- lations.” Studies on Neotropical Fauna and Environment 47 (1): 19–23. doi:10.1080/01650521.2012.662795 .

Roberts, Gilbert. 1996. “Why Individual Vigilance Declines as Group Size In- creases.” Animal Behaviour 51 (5): 1077–86. doi:10.1006/anbe.1996.0109.

Slabbekoorn, Hans, and Ardie den Boer-Visser. 2006. “Cities Change the Songs of Birds.” Current Biology 16 (23): 2326–31. doi:10.1016/j.cub.2006.10.008.

Tan, Ashley. 2019. “Monitor Lizard Steals Huge Fish Belonging to Otter Family at Botanic Gardens.” Mothership.sg. https://mothership.sg/2019/08/monitor-lizard-steal-fish-otter-zouk5/.

Tan, Mei Mei. 2017. “Behavioural Patterns of Intergroup Territorial Agonism in Lutrogale Perspicillata in Singapore.” Undergraduate Thesis, Yale-NUS College.

Team, RStudio. 2016. “RStudio: Integrated Development Environment for R.” Boston, MA: RStudio Inc. http://www.rstudio.com/.

Theng, Meryl, and N Sivasothi. 2016. “The Smooth-Coated Otter Lutrogale Perspicillata (Mammalia: Mustelidae) in Singapore: Establishment and Expansion in Natural and Semi-Urban Environments.” IUCN/SCC Otter Specialist Group Bulletin 33 (January): 37–49.

Theng, Meryl, Sivasothi N., and Heok Tan. 2016. “Diet of the Smooth-Coated Otter Lutrogale Perspicillata (Geoffroy, 1826) at Natural and Modified Sites in Singapore.” The Raffles Bulletin of Zoology 64 (September): 290–301.

Toh, Ting Wei. 2018. “Scuffle Between Crocodile and Otters Captured on Video in Sungei Buloh Not a Rare Sight: Nature Observers, Environment News & Top Stories - the Straits Times.” Newspaper. The Straits Times. https://www.straitstimes.com/singapore/environment/scuffle-between-crocodile-and-otters-captured-on-video-not-a-rare-sight-nature

Tosa MI, Dziedzic EH, Appel CL, Urbina J, Massey A, Ruprecht J, Eriksson CE, Dolliver JE, Lesmeister DB, Betts MG, Peres CA, Levi T (2021) The rapid rise of next-generation natural history. Front Ecol Evol 9:698131. https://doi.org/10.3389/fevo.2021.698131

Twining, Joshua P, and André Koch. 2018. “Dietary Notes and Foraging Ecology of South-East Asian Water Monitors (Varanus Salvator) in Sabah, Northern Borneo, Malaysia.” The Herpetelogical Bulletin 143: 36–38.

Uyeda, Linda. 2009. “Garbage Appeal: Relative Abundance of Water Monitor Lizards (Varanus Salvator) Correlates with Presence of Human Food Leftovers on Tinjil Island, Indonesia.” Biawak 3 (1): 9.

Wickham, Hadley, and RStudio. 2019. “Tidyverse: Easily Install and Load the ‘Tidyverse’.” https://CRAN.R-project.org/package=tidyverse.

Wong, George. 2019. RandomSG. YouTube. Accessed 23 June 2022. www.youtube.com/channel/UCLz7pIXxzaFzz_MN02kNzsQ

